# A high-throughput method for quantifying *Drosophila* fecundity

**DOI:** 10.1101/2024.03.27.587093

**Authors:** Andreana Gomez, Sergio Gonzalez, Ashwini Oke, Jiayu Luo, Johnny B. Duong, Raymond M. Esquerra, Thomas Zimmerman, Sara Capponi, Jennifer C. Fung, Todd G. Nystul

## Abstract

Measurements of Drosophila fecundity are used in a wide variety of studies, such as investigations of stem cell biology, nutrition, behavior, and toxicology. In addition, because fecundity assays are performed on live flies, they are suitable for longitudinal studies such as investigations of aging or prolonged chemical exposure. However, standard Drosophila fecundity assays have been difficult to perform in a high-throughput manner because experimental factors such as the physiological state of the flies and environmental cues must be carefully controlled to achieve consistent results. In addition, exposing flies to a large number of different experimental conditions (such as chemical additives in the diet) and manually counting the number of eggs laid to determine the impact on fecundity is time-consuming. We have overcome these challenges by combining a new multiwell fly culture strategy with a novel 3D-printed fly transfer device to rapidly and accurately transfer flies from one plate to another; the RoboCam, a low-cost, custom built robotic camera to capture images of the wells automatically; and an image segmentation pipeline to automatically identify and quantify eggs. We show that this method is compatible with robust and consistent egg laying throughout the assay period; and demonstrate that the automated pipeline for quantifying fecundity is very accurate (r^2^ = 0.98 for the correlation between the automated egg counts and the ground truth) In addition, we show that this method can be used to efficiently detect the effects on fecundity induced by dietary exposure to chemicals. Taken together, this strategy substantially increases the efficiency and reproducibility of high throughput egg laying assays that require exposing flies to multiple different media conditions.

## Introduction

Measurements of Drosophila fecundity are used in a wide variety of studies, including investigations of aging, stem cell biology, nutrition, behavior, and toxicology [1–7]. These studies build on the wealth of knowledge about Drosophila oogenesis and the complex array of inputs that combine to optimize the rate of egg laying in response to the environment. Environmental cues are sensed primarily through pheromones [8–10], which are detected by the olfactory system, and nutrient cues, which are received through the digestive tract and sensed by the fat body [11–13]. These upstream signals coordinate the release of hormonal signals into the hemolymph that regulate the cell and tissue level responses throughout the body. For example, within the ovary, these signals act on both the stem cell compartment, called the germarium, to regulate the rate of cell division [14–16] as well as on later stages of oogenesis to promote follicle survival and maturation [17–19]. Intercellular signals also trigger ovulation of mature eggs into the uterus, fertilization, and egg deposition [20–23]. Thus, measurements of fecundity provide a real-time, quantitative assessment of this multiorgan process in live flies. This makes the assay suitable for longitudinal studies such as investigating how fecundity changes with age or with prolonged chemical exposure. However, as the culmination of a broad cascade of inputs, fecundity assays must be carefully designed to ensure robust and reproducible results.

Robust egg laying occurs during the first four weeks of adult life [1]. Environmental conditions that promote a high, consistent rate of egg laying include the presence of males, a lack of overcrowding, room temperature, and a protein rich diet, which is usually provided by the addition of a wet yeast paste to culture vials with standard media [11,24]. Under these conditions, each female typically lays approximately 60-100 eggs per day [25]. A common approach to quantifying the rate of egg laying is to maintain flies in standard culture vials or molasses agar plates for a defined period of time and then to manually count the number of eggs that are visible under a dissecting microscope [23,25,26]. This method is simple and has the advantage that flies are maintained in standard conditions, but it is time consuming and prone to variation due to human error across observations. In addition, this approach may underestimate the rate of egg laying because agar-based culture media and yeast paste are soft, so eggs can sink below the surface and become obscured from view.

Therefore, several studies have modified this approach to address these challenges. For example, using a firmer agarose-based culture media is useful for keeping eggs on the surface [27], though eggs deposited directly in the yeast paste (if it is added) are still hard to see. To increase speed and improve reproducibility, several studies have developed methods to automate the process of identifying and counting eggs. These approaches start with images of eggs laid in a standard culture vial or large plate and apply an image segmentation algorithm to automatically identify and quantify the eggs [28–30]. Lastly, other studies have used multiwell plates rather than culture vials to increase the efficiency and reduce the cost of testing many different diets or chemical exposures [31–33]. However, a major drawback to the multiwell format is the difficulty of ensuring that flies remain in the same wells throughout the assay and of accurately transferring flies from one plate to another, as is typically needed for a multi-day time course. Each of these innovations is useful on its own but they have been optimized for a particular application and thus are not easily combined into a single workflow.

Here, we describe a strategy for performing a high throughput multi-day fecundity assay and demonstrate that it is effective for measuring the impact of chemical additives in the diet. Building on previous innovations, our strategy combines a multiwell format with a novel 3D- printed fly transfer device to rapidly and accurately transfer flies from one plate to another; the RoboCam, a low-cost, custom built robotic camera to automatically capture images of the wells; and an image segmentation pipeline to automatically identify and quantify eggs. We show that this method is compatible with robust and consistent egg laying throughout the assay period and that it can accurately detect chemical-induced effects on fecundity. Taken together, this strategy substantially increases the efficiency and reproducibility of high throughput egg laying assays that require exposing flies to multiple different media conditions.

## Methods

### Culturing conditions and fecundity assay

The egg laying medium is made by diluting 100% Concord Grape Juice (Santa Cruz Organic) in water to a final concentration of 20% grape juice and adding sucrose (Sigma-Aldrich, 84100) at 0.2 g/ml and agarose powder (Genesee Scientific, 20101) at 0.02 g/ml. The solution is boiled to melt the agarose powder, and 300 µL of the hot solution is added to each well, with care taken to minimize bubbles so that the surface is flat. Once cooled, 30 µL of a wet yeast slurry (0.2 g/mL of Red Star yeast granules dissolved in water) is added on top of the media and the sides of the plate are tapped until the solution evenly covers the entire surface of the gel. The plates are then dried for in a fume hood until the yeast layer is completely dry (approximately 90 minutes). This is important to ensure that the flies do not get stuck in the media.

Flies from the *Drosophila melanogaster* strain, *w1118* (BIDC Stock # 3605) are raised on standard molasses food at 25°C. The bottles are cleared of adults and then newly eclosed flies are fed wet yeast daily for one week before starting the fecundity assays. Flies are then maintained on an egg laying medium with a layer of wet yeast on top in 48-well cell culture plates (Genesee Scientific, 25-103) for a 24-hour “pre-egg laying period.” This provides time for the small cohorts of flies in each well to become accustomed to the new environment and receive a uniform nutritional experience before the fecundity measurements begin.

After the pre-egg laying period, the fecundity assay begins. Flies are maintained on the egg laying medium with yeast on top as described above, either without any other additives, with DMSO, or with a chemical dissolved DMSO, as described in the “Preparation of chemical solutions” section below. Flies are transferred to new plates containing fresh medium with yeast on top every day during days when quantification of fecundity is not required. On days when quantification of fecundity is required, the yeast slurry is added to the egg laying medium while it is still warm, and the solution is mixed to homogeneity before pipetting into the wells. For chemical exposure conditions, the chemical solution is added on top after the medium hardens and dried in a fume hood, as described above. This modification is important to provide an optically uniform surface for egg laying while still providing flies with adequate nutrition and chemical exposure.

### Construction of the 3D-printed fly transfer device

The fly transfer lid was constructed from a sheet of 0.6 mm nylon mesh mounted between a lower array of cups (8mm diameter, 10 mm deep, 6-degree taper) and an upper array of gas exchange holes. The lower and upper layers were designed with Tinkercad (www.tinkercad.com) and stereo lithographically 3D printed with LEDO 6060 resin (www.jlcpcb.com). Studs (3 mm tall, 2.5 mm diameter) in the upper layer fit into matching holes (3 mm diameter) in the nylon sheet and lower layer, sealed together with cyanoacrylate glue. The holes in the nylon mesh were formed by melting the desired nylon hole locations with a hot soldering iron guided by a metal template.

### Adding and transferring flies using a 3D-printed fly transfer device

A 3D-printed transfer device is used to keep the flies in individual wells during the culture period and ensure accurate transfer to new wells. The transfer device is placed on a CO2 pad and two females and one male are added into each cylinder of the transfer device. The plate containing the egg laying medium is placed on top of the lid and secured to ensure proper fit into each well. The plate and transfer device are removed from the CO2 pad and flipped over so that the flies land on the media. When the flies recover and begin flying, the plate, with the transfer device as a lid, is placed in a 25°C incubator. To transfer flies from one plate to another, the plate and transfer device is placed mesh-side down on a CO2 pad. Once the flies have dropped to the mesh lining of the lid, the current plate is removed and replaced with a new plate. The older plate is either saved for imaging or discarded.

### Construction of the RoboCam

To build the RoboCam a high-resolution digital camera (12 MP HQ Camera, Adafruit) equipped with a 16mm telephoto lens (Adafruit) is attached to the head of a conventional 3D printer (AnyCubic Mega S) and an LED Light Tracing Box is placed on the 3D printer plate, so that once the well plates are positioned, it illuminates them from below (**Figure S1**). A single-board, Raspberry Pi computer (Adafruit) controls the x, y, and z camera location using G-code commands. G-code is a standard programming language for 3D printer that provides instructions to the machine on where to move, how fast, and which path to follow. The G-code commands are generated by the user with a Graphical User Interface (GUI) running on the Raspberry Pi, written in Python with PySimpleGUI and OpenCV libraries. The well images collected by the RoboCam are stored on an external hard drive.

### Imaging

To collect egg laying data, flies are removed exactly 24 hours after placement into the wells and the plate is either placed in a -20°C freezer for imaging later (no more than 48 hours) or imaged immediately. For imaging, the plate is placed on the RoboCam platform; the camera position is registered to the wells in each of the four corners; the z-focus is adjusted as needed; and the movement and acquisition module is initiated. This module moves the camera to a precise position over each well and takes a single image in the specified z-position. If this z-position is not optimal for some wells, the module can be run again at another z-position.

### Imaging pipeline implementation

The imaging pipeline is written in Python and provided as a Google CoLab notebook. To run the imaging pipeline, the necessary input and output file paths and the estimated well diameter are provided as inputs. Then, Stardist [34,35] is installed (https://github.com/stardist/stardist), the necessary libraries are loaded, several functions are defined, and the imaging processing steps are performed. Briefly, the image processing steps are (1) Preprocessing, which includes inverting the image, converting it to 8-bit grey scale, and normalizing the pixel intensity; (2) Well detection using the canny edge detection module from the Python package, skimage; (3) Image segmentation using flyModel2 (the default probability threshold of 0.7 can be manually adjusted based on the quality of the images) to identify eggs; (4) A filtering step, in which segmented objects are that are outside of the predicted well, less than 2500 pixels^2^, or greater than 4000 pixels^2^ are excluded; and (5) an Output step in which the number of objects (eggs) per well are exported as an Excel file and images showing the segmentation results are saved.

### Quantifly analysis

Quantifly [30] was trained using 19 images according to the instructions provided by the manual. The software was trained at different values of sigma. Sigma value of 2 gave the most accurate counts and was used for comparison with the Stardist models.

### Stardist model training

52 randomly chosen images were used for annotation using Labkit plugin in Fiji. The resulting pairs of images and masks were used to train a Stardist model by adapting the example Python notebook (https://github.com/stardist/stardist/blob/master/examples/2D/2_training.ipynb) for this purpose. Data augmentation and training was performed as suggested with a split of 44 training images and 8 validation images. The optimized probability threshold for flyModel2 is 0.706 and the nms_thresh is 0.3. The key metrics for flyModel2 at t=0.5 are listed: fp = 3; tp = 50; fn = 2; precision = 0.9434; recall = 0.9615; accuracy = 0.909.

### Preparation of chemical solutions

1000x stock solutions in 100% DMSO (Sigma-Aldrich, D8418) are diluted 1:100 with water to achieve a 10x concentration of chemical in 1% DMSO. Then, 30 µL of the 10x chemical solution is added into each well to achieve a final concentration of 1x chemical in 0.1% DMSO. Specifically, to achieve 0.1 µM, 10µM and 25 µM final concentrations of rapamycin (bioWORLD, 41810000-2) and bendiocarb (Sigma-Aldrich, 45336), the 1000x stock solutions were 0.1 mM, 10 mM, and 25 mM, respectively.

### Chemical exposure time course

To quantify the effect of chemical exposure on fecundity and viability, flies are maintained in the 48-well plates with or without chemical, transferred to a fresh plate every day, for 7 days. The egg laying medium with the yeast mixed in is used on days 1, 3 and 7 so that fecundity can be quantified and egg laying medium with the yeast on top is used on all other days. To thoroughly assess reproducibility, each chemical condition was repeated in at least 16 wells per replicate, and data are an aggregate of seven replicates. The seven replicates do not all include every condition and time point, but there were at least 60 wells total across all replicates for every condition and time point (see FigureS2.Rmd in the Github repository for more details). Fecundity data was not collected from any condition in which one or more females do not survive in at least half of the wells. Otherwise, egg laying counts are performed on all wells in which both females are alive whereas the data from wells with fewer than two females alive are discarded.

### Data and code availability

All raw data and software code are publicly available on the Nystul Lab Github site at https://github.com/NystulLab/HighThroughputFecundityAssay except the 3D stereolithography (STL) design files, which are publicly available on the Center for Cellular Construction Github site at https://github.com/CCCofficial/FlyTransferCup48.

## Results

### Multi-well culture conditions

The first step in creating a high-throughput assay for quantifying fecundity was to develop a procedure for culturing flies in a 48-well format that promoted robust, reproducible egg laying onto a surface that could be clearly imaged and is compatible with controlled exposure to chemicals or other additives to the diet. We found that a standard grape juice and agarose recipe is a good base, as it is firm enough to prevent eggs from sinking below the surface, optically uniform, and resistant to drying and cracking over a 24-hour period. However, the typical procedure of adding a dollop of yeast paste to the side of the chamber is too cumbersome and imprecise for the high-throughput 48-well format. In addition, with the yeast paste physically separated from the base media, the nutritional experiences and exposures to chemicals in the diet are more variable as they depend on the amount that individual flies consume from each nutritional source. Mixing dried yeast into the grape juice and agarose mixture before it hardened is more efficient and ensures a uniform nutritional experience. However, while this method supports robust egg laying for 24 hours, we found that fecundity drops significantly when flies are maintained on this media for more than one day (**Figure 1**). This is likely due to an absence of wet yeast paste, rather than the composition of the media, as similar results were obtained when we used the standard molasses or BIDC recipes (**Figure 1**).

**Figure 1:**
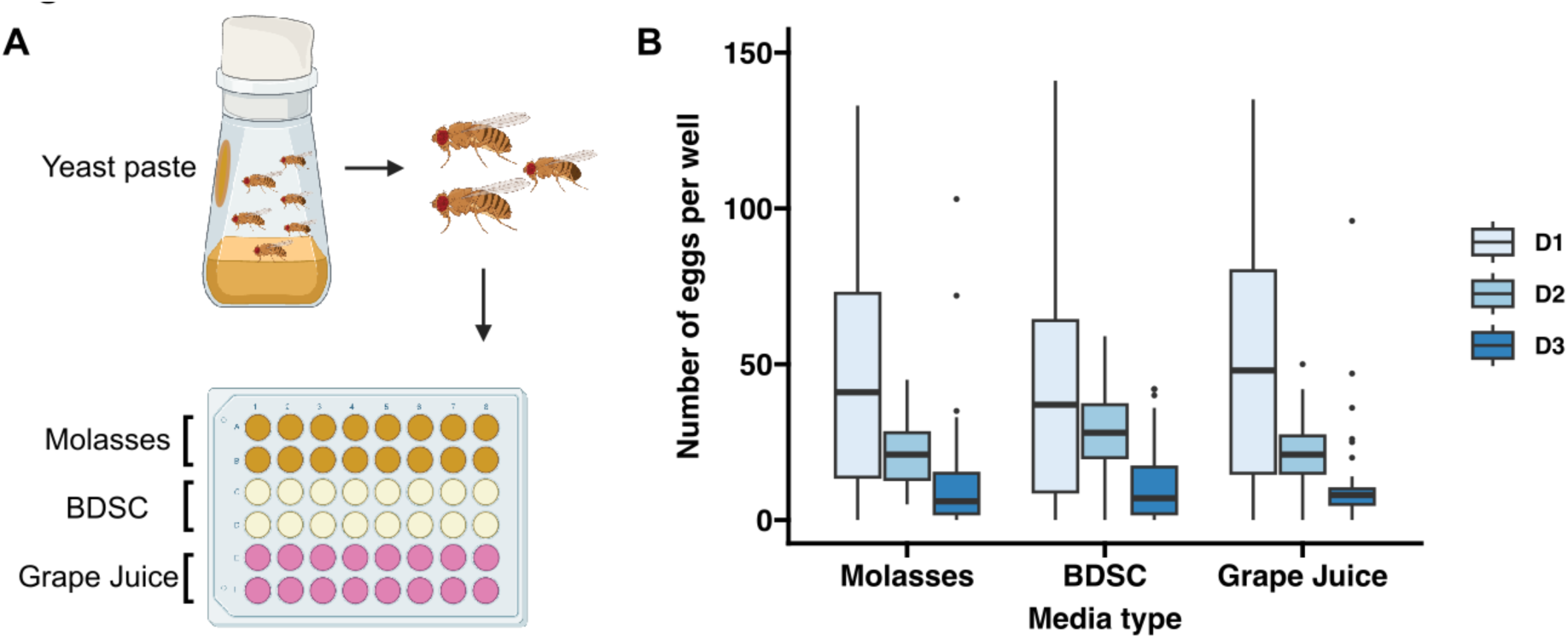
Quantification of fecundity over three days on different types of media. (A) Diagram showing the workflow for this assay. Flies were fed wet yeast paste for two consecutive days and then 2 females and one 1 male were transferred into each well of a 48- well plate with molasses-based, BDSC-based, grape juice-agarose based media (16 wells for each condition) and allowed to lay eggs for 23 hours. (B) Graph showing the number of eggs laid in each condition, as determined by manual counts.

We therefore investigated a third option, which was to apply a defined amount of a wet yeast slurry (created by adding more water than would be used to make yeast paste) to the surface of the media with a pipetman and then allowing the excess water to evaporate from the plates so that a moist, even layer of yeast coated the entire surface of the media. This created a uniform dietary exposure and supported robust egg laying but we found that it is difficult to reliably detect all the eggs laid in this condition because the yeast layer is soft and not optically uniform.

Therefore, we devised a protocol that took advantage of all three of these methods (**Figure 2A**). First, a large cohort of freshly eclosed flies are maintained in standard culture bottles with a dollop of wet yeast added to the side of the chamber. This allows the flies to reach sexual maturity with a standard, protein-rich diet. Then, two females and one male are transferred into each well of a 48-well plate that has been prepared with the layer of wet yeast on top, as described above. This number of flies was chosen because we found that there was lower interwell variability with two females compared to one while more than three flies per well increased the frequency of wells that needed to be excluded because one or more flies in the well died during the assay period. After one day of acclimating to the environment in the 48-well plates, the flies are transferred to a new 48-well plate with the wet yeast layer on top every other day so they have constant access to a moist, protein rich diet that exposes them to the desired dietary condition. To assess fecundity, the flies are transferred to a new 48-well plate that has been prepared with the yeast mixed into the base media, as described above. After 24 hours, the flies are transferred back to a plate with yeast on top if the time course is continuing or discarded if not, and the eggs are imaged and quantified, as described below. We found that alternating in this way between the condition with the wet yeast on top and the condition with the yeast mixed into the media at time points when fecundity is measured was still able to support a consistent rate of egg laying for at least 3 days, with only a modest drop off by 7 days (**Figure 2B-E**). Specifically, we observed an average of 35.9, 42.7, and 19.8 eggs per well after 1, 3, and 7 days, respectively. While this is lower than the 60-100 eggs per female that has been reported in some studies using soft media [25], it is consistent with a previous finding that the rate of egg laying decreases with increasing stiffness of the media [27]. Furthermore, as we demonstrate below, the number of eggs per well is high enough that dose-dependent effects of chemical exposure could still be detected. Thus, we find this protocol to be a useful compromise that balances physiological needs with accuracy and efficiency.

**Figure 2:**
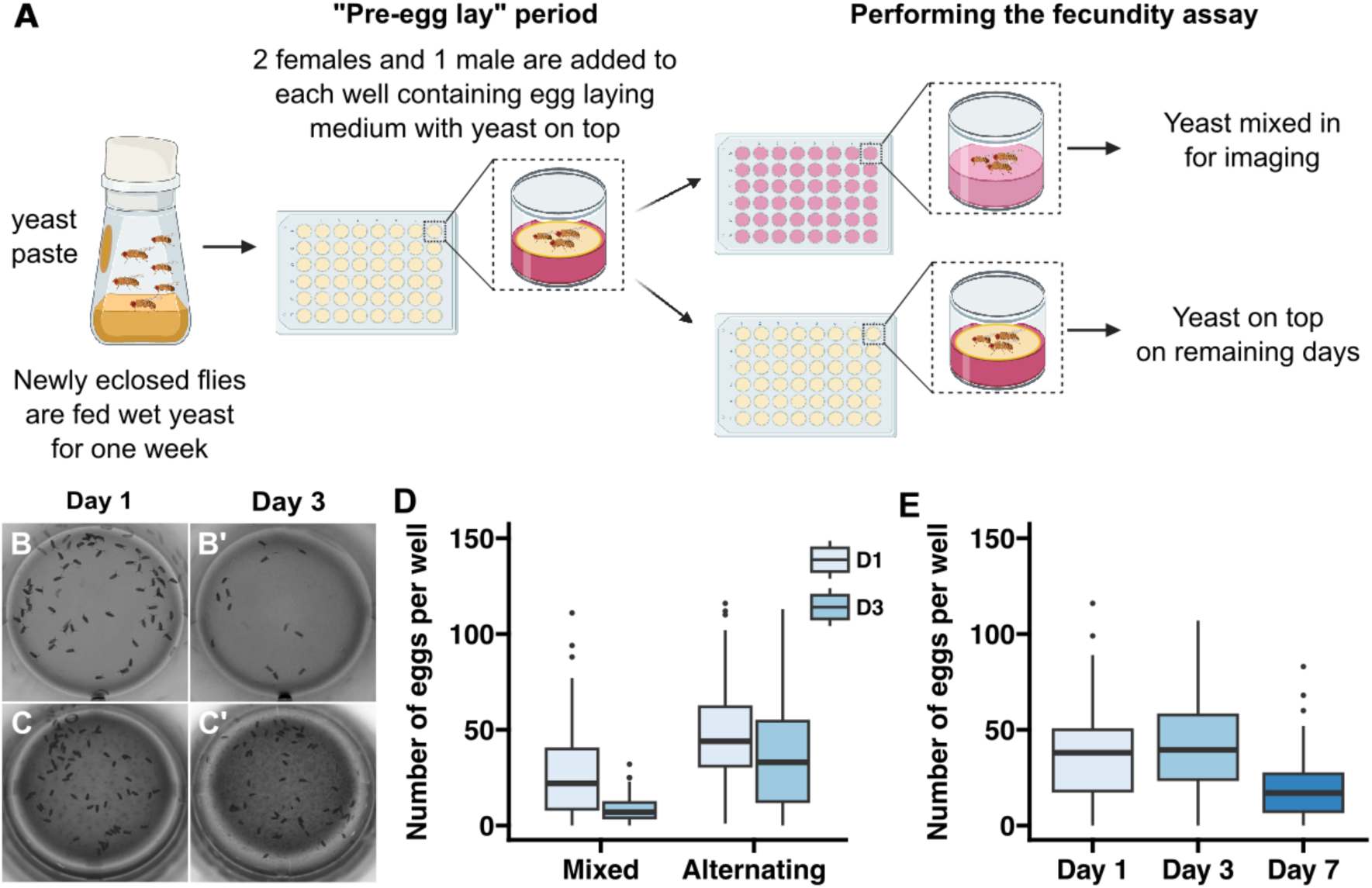
Development of protocol for assaying fecundity over 7 days. (A) Diagram showing a workflow in which flies are maintained on media with yeast mixed in on the days in which images will be acquired to assess fecundity and on media with yeast on top during the remaining days. In the “alternating” conditions shown in panels C-E, flies were put on media with yeast mixed in on days 1, 3, and 7 and on media with yeast on top on days 2, 4, 5, and 6. (B-D) Representative examples of wells from a 3 day time course in which flies were maintained on wells with the yeast mixed into the media at all time points (B) or alternating between wells with yeast mixed into the media on days 1 and 3 and yeast on top on day 2 (C), and a graph showing the number of eggs laid in each regime (D). (E) Graph showing the number of eggs laid in the alternating regime over a 7-day time course.

### Fly transfer device

The 48-well plate format is useful for testing multiple different experimental conditions in a high- throughput manner. However, the plastic lid does not fit tightly enough over the surface of the wells to ensure that flies cannot escape or move between wells, and it is very difficult to anesthetize and transfer the flies to a new plate without losing some or mixing them up. In addition, manually transferring each fly to the new well is a time-consuming process. To overcome these challenges, we designed a custom transfer device with tapered cylinders that fit snugly into each well of a 48-well plate (**Figure 3**). The wide ends of the cylinders are blocked off with a fine mesh that allows for air exchange while preventing the flies from escaping. To transfer the flies to a new 48 well plate, the flies are anesthetized by inverting the transfer device and plate onto a CO2 pad so that the flies fall onto the mesh surface and the old plate can be replaced with a new one. The transfer device and new plate are removed from the CO2 pad and, when the flies recover, the transfer device and plate are inverted so that the plate is upright again. This process is efficient, reducing the transfer time from approximately 5 minutes to less than 30 seconds, and ensures that flies are accurately transferred to the new wells and are unable to escape during the incubation periods. An additional benefit is that flies are only exposed to CO2 for a very brief period of time (less than a minute) so the potential effects that CO2 may have on the health or fecundity of the flies is minimized.

**Figure 3:**
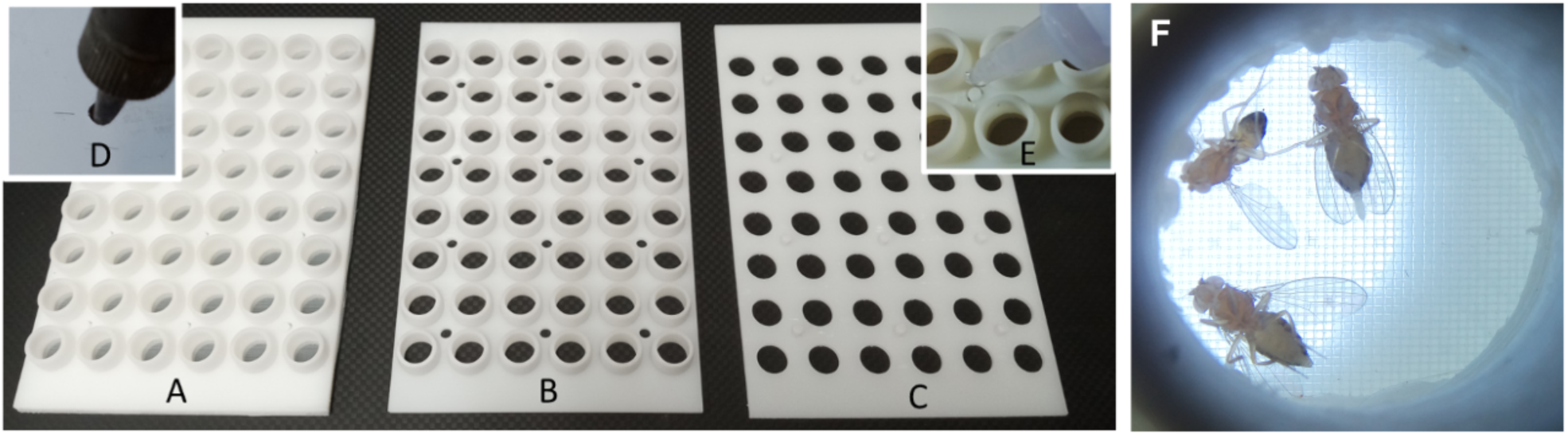
Fly transfer device. A fly transfer lid (A) is constructed from a layer of nylon mesh bonded between an array of cups (B) and holes (C). Studs in the hole layer (C) pass through corresponding holes in the cup (B) and nylon layers, creating individual gas exchange tops for each well. A hot soldering iron, guided by holes in a metal template, creates holes in the nylon mesh (D). The final assembly (E) studs are bonded with cyanoacrylate glue. When placed on a CO2 plate, anesthetized flies fall into their corresponding cup, enabling the plate to be replaced and flipped, simultaneously transferring all flies into new wells.

### RoboCam platform

Our next goal was to develop a high-throughput method to i_A_mage the surface of each well. We first experimented with different forms of illumination and found that lighting from underneath using an ultra-thin LED Light Tracing Box was ideal, as it minimized shadows and reflections from the plastic and provided a good contrast between the eggs and the media. Then, we developed the RoboCam platform to automate the image acquisition. This platform integrates the precise robotic functionality of a 3D printer with the computational efficiency of Raspberry Pi computers and a high-resolution digital camera with a telephoto lens into a programmable robotic camera (**Figure 4A-G** and **Table 1**). An external hard drive connected to the Raspberry Pi ensures a safe storage of the images. We developed a Graphical User Interface (GUI) in Python to control the system (**Figure S1**). Through this interface, the Raspberry Pi issues serial commands to control the RoboCam, facilitating precise movements in a snake path across the plate to sequentially position the camera over each well (**Figure 4H**), capture images, and store the image files on the external hard drive. Before each run, the RoboCam is calibrated to the well plate by aligning a circular target with the wells in each of the four corners of the plate, which is stored as a CSV file along with the well scan pattern. Focus is manually set on the lens or by moving the camera in the z direction through the GUI. Upon completion of the run, the images stored on the external hard drive are transferred to another computer for the image analysis steps. The advantages of using the RoboCam platform to collect images are discussed in detail in the Discussion section. Scientifically, the main benefit of the RoboCam is the consistency of the images. In all the images taken by the RoboCam, the wells are in the exact same position, greatly facilitating the data analysis. On the technical side, the modular set-up of the RoboCam allows changes to its design, so that scientists can modify it depending on the specific purpose of their research.

**Figure 4:**
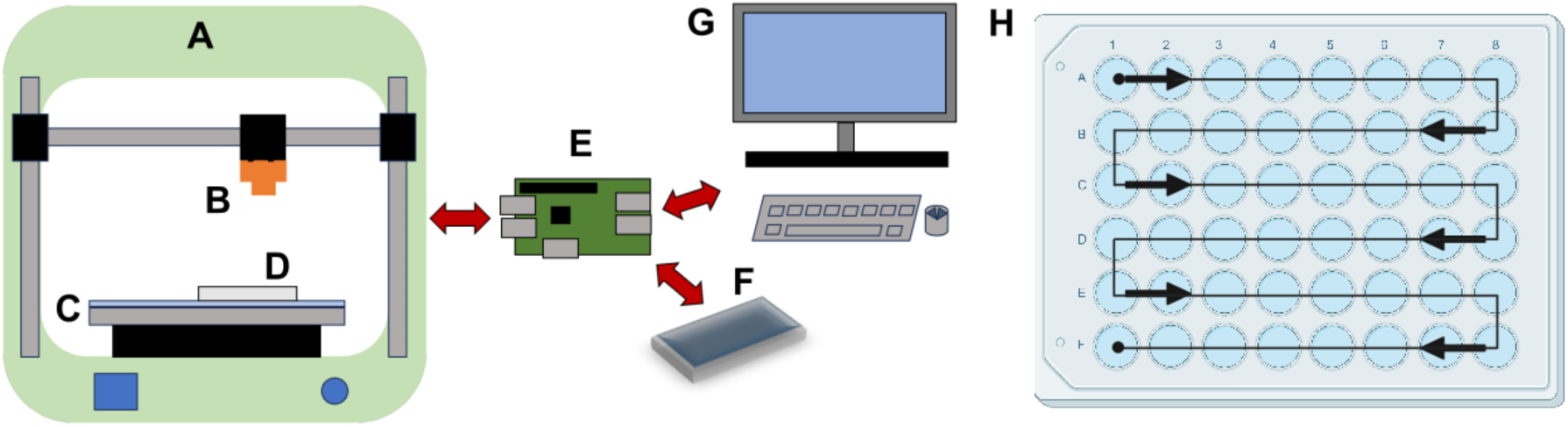
RoboCam device for automated image capture. A 3D printer (A) is modified by adding a camera (B) and light plate (C), where the 48-well plate (D) is located. The light ensures good contrast between the eggs and the media, while minimizing shadows and reflections. A single-board computer (E) controls the x, y, and z movement of the camera (B), and saves captured images on a hard drive (F). The user programs the RoboCam using a graphical user interface with a keyboard, mouse, and monitor (G). (H) Camera system moves in a snake path, centering over individual wells.

**Table 1.**
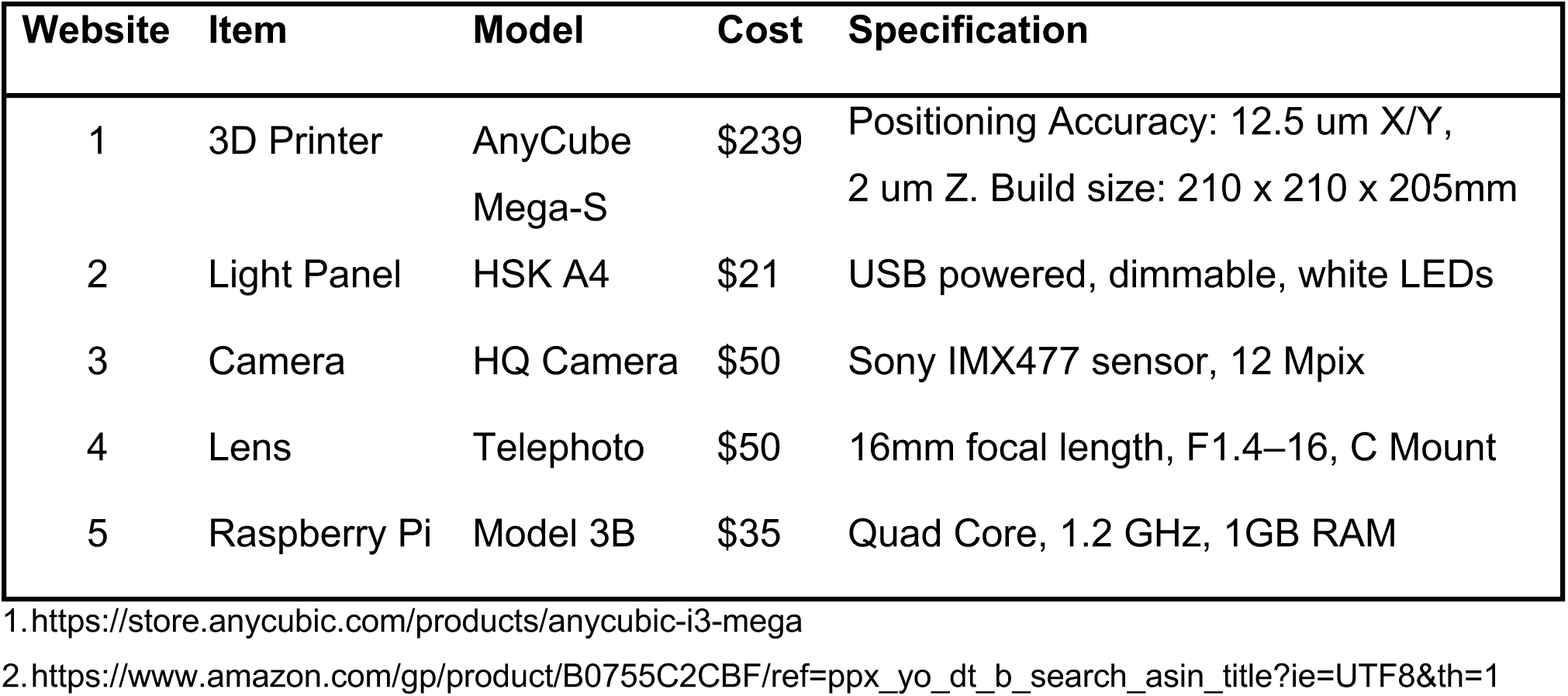
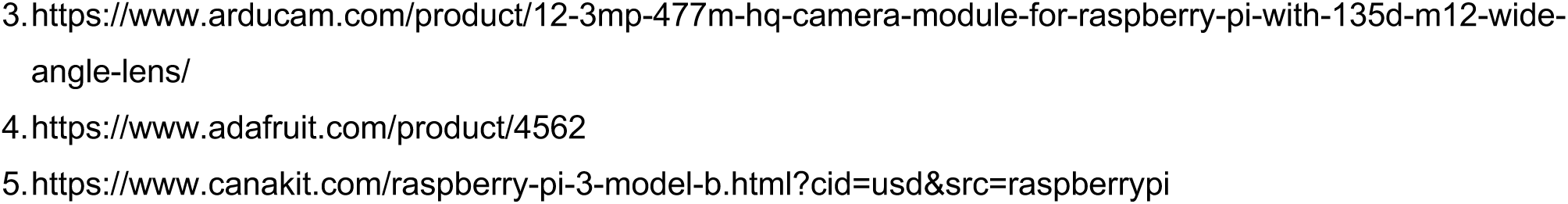
RoboCam Components.

| Website | Item | Model | Cost | Specification |
| --- | --- | --- | --- | --- |
| 1 | 3D Printer | AnyCube<br>Mega-S | \$239 | Positioning Accuracy: 12.5 μm X/Y,<br>2 μm Z. Build size: 210 x 210 x 205mm |
| 2 | Light Panel | HSK A4 | \$21 | USB powered, dimmable, white LEDs |
| 3 | Camera | HQ Camera | \$50 | Sony IMX477 sensor, 12 Mpix |
| 4 | Lens | Telephoto | \$50 | 16mm focal length, F1.4–16, C Mount |
| 5 | Raspberry Pi | Model 3B | \$35 | Quad Core, 1.2 GHz, 1GB RAM |

1. https://store.anycubic.com/products/anycubic-i3-mega
2. https://www.amazon.com/gp/product/B0755C2CBF/ref=ppx_yo_dt_b_search_asin_title?ie=UTF8&th=1
- 1 3. [https://www.arducam.com/product/12-3mp-477m-hq-camera-module-for-raspberry-pi-with-135d-m12-wide-](https://www.arducam.com/product/12-3mp-477m-hq-camera-module-for-raspberry-pi-with-135d-m12-wide-angle-lens/) - 2 [angle-lens/](https://www.arducam.com/product/12-3mp-477m-hq-camera-module-for-raspberry-pi-with-135d-m12-wide-angle-lens/) - 3 4. https://www.adafruit.com/product/4562 - 4 5. https://www.canakit.com/raspberry-pi-3-model-b.html?cid=usd&src=raspberrypi

### Image segmentation

With the steps described above in place, we next sought to develop an image segmentation pipeline that would automatically identify and quantify eggs from the images taken by the RoboCam. First, we tested Quantifly [30], which utilizes a density estimation approach to quantify the number of dense objects in the image. To determine the accuracy of this approach, we carefully hand-counted the number of eggs in 133 images and compared these manual counts to the results from Quantifly using multiple different sigma values. The best performance we could obtain from Quantifly on this dataset was an R^2^ = 0.74 (**Figure 5A, D**). Next, we investigated whether Stardist, a deep learning method that is based on star-convex shape detection [34,35], could achieve better performance. Indeed, the pretrained Stardist model, 2D_versatile_fluo, achieved an of R^2^ = 0.80 with the same set of images (**Figure 5B, D**). To improve upon these results, we generated a custom model (flyModel2) trained on a set of images with hand-drawn outlines around each egg as ground-truth annotations. This increased the accuracy to an R^2^ = 0.91. We examined the sources of the errors in this approach and noticed that, in some cases, eggs near the edge of the well were reflected in the clear plastic wall and the reflections were counted as separate eggs (**Figure 5E**). Increasing the sensitivity threshold reduced the number of these false positives but also increased the number of false negatives. Thus, to eliminate this type of error, we used an ellipse detection tool to identify the well boundary and restrict the analysis to the portion of the image within this boundary (**Figure 5F**). The RoboCam positions each well in precisely the same location in each image, so we were able to accurately map the well in all the images from the same plate by performing the well detection on the first well and then using the coordinates to position the mask thereafter (**Figure 5G**). Together these strategies comprise a highly accurate (R^2^ = 0.98) and efficient pipeline for quantifying egg number (**Figure 5C-D**).

**Figure 5:**
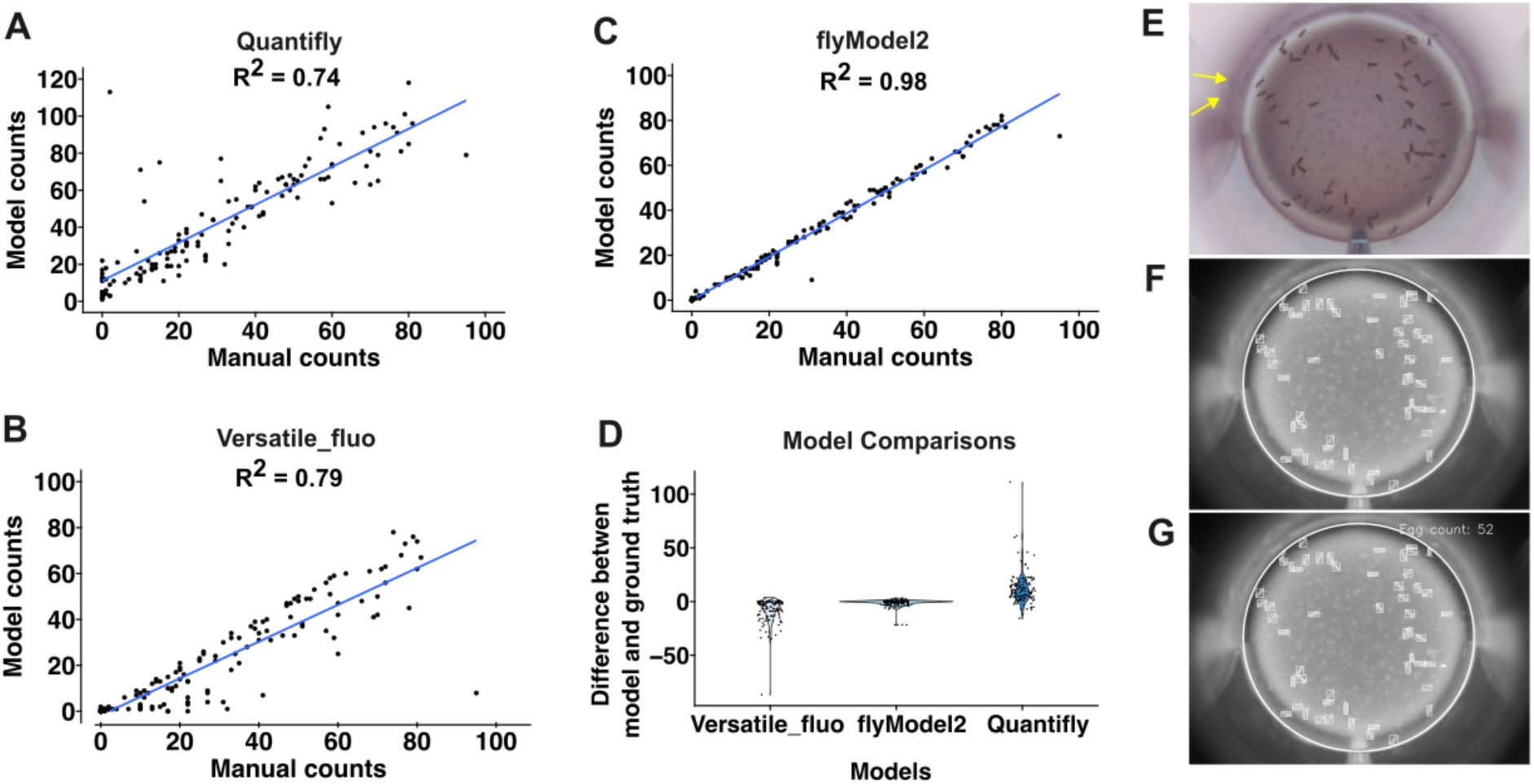
Automated image analysis pipeline. (A-C) Graphs showing comparisons of manual egg counts to automated egg counts using Quantifly (A), Stardist with the Versatile_fluo model (B), or Stardist using a custom-build model, flyModel2 (C). Each dot shows the manual and automated counts from a single well. (D) Graph showing the error between manual and automated egg counts. Each dot is the difference between the automated count and the manual count (ground truth). (E-G) Images showing individual steps in the automated image analysis pipeline. Starting from the raw image (E), the Stardist model identifies the eggs and the elipse detection tool identifies the well edge (boxed regions and white circle, respectiely in Panel F). Then, the number of boxed regions (eggs) within the well are counted (G). Eggs reflected in the well walls are indicated with yellow arrows.

### Quantification of the effects of chemical exposure

To evaluate the performance of this pipeline in experimental contexts that alter fecundity, we used it to quantify egg laying over a 7-day time course of exposure to either rapamycin, which is a potent inhibitor of Tor signaling that reduces fecundity in Drosophila [36], or and bendiocarb, which is an insecticide used to control mosquito populations [37] (**Figure 6A**). Both chemicals have been shown to have effects in the micromolar range [36,38], so we tested 0.1 μM, 10 μM, and 25 μM doses. For both chemicals, we observed a progressive decrease in the number of eggs with increased dose and length of exposure (**Figure 6B-F**). However, these observations are based on quantification of eggs from at least 60 wells per condition. To assess whether the effects of rapamycin and bendiocarb in the diet could have been detected with fewer repeats of each condition, we downsampled the data by randomly choosing 4, 8, 16, or 24 wells from each condition (day and chemical dose) 100 times and performing pairwise T-tests on each of these randomly selected subsets of the data (**Figure S2**). We found that nearly all of tests with 16 or 24 wells sampled from the Day 3 and Day 7 datasets reproduce the conclusions about statistical significance from the entire dataset (except for 0.1 µM bendiocarb at Day 7) and that most tests with 8 wells sampled were able to detect a statistically significant difference in fecundity at the highest (25 µM) dose of rapamycin and bendiocarb at Day 3 (**Figure 6**). Thus, this pipeline is able to efficiently detect dose and time dependent experimentally induced decreases in fecundity.

**Figure 6:**
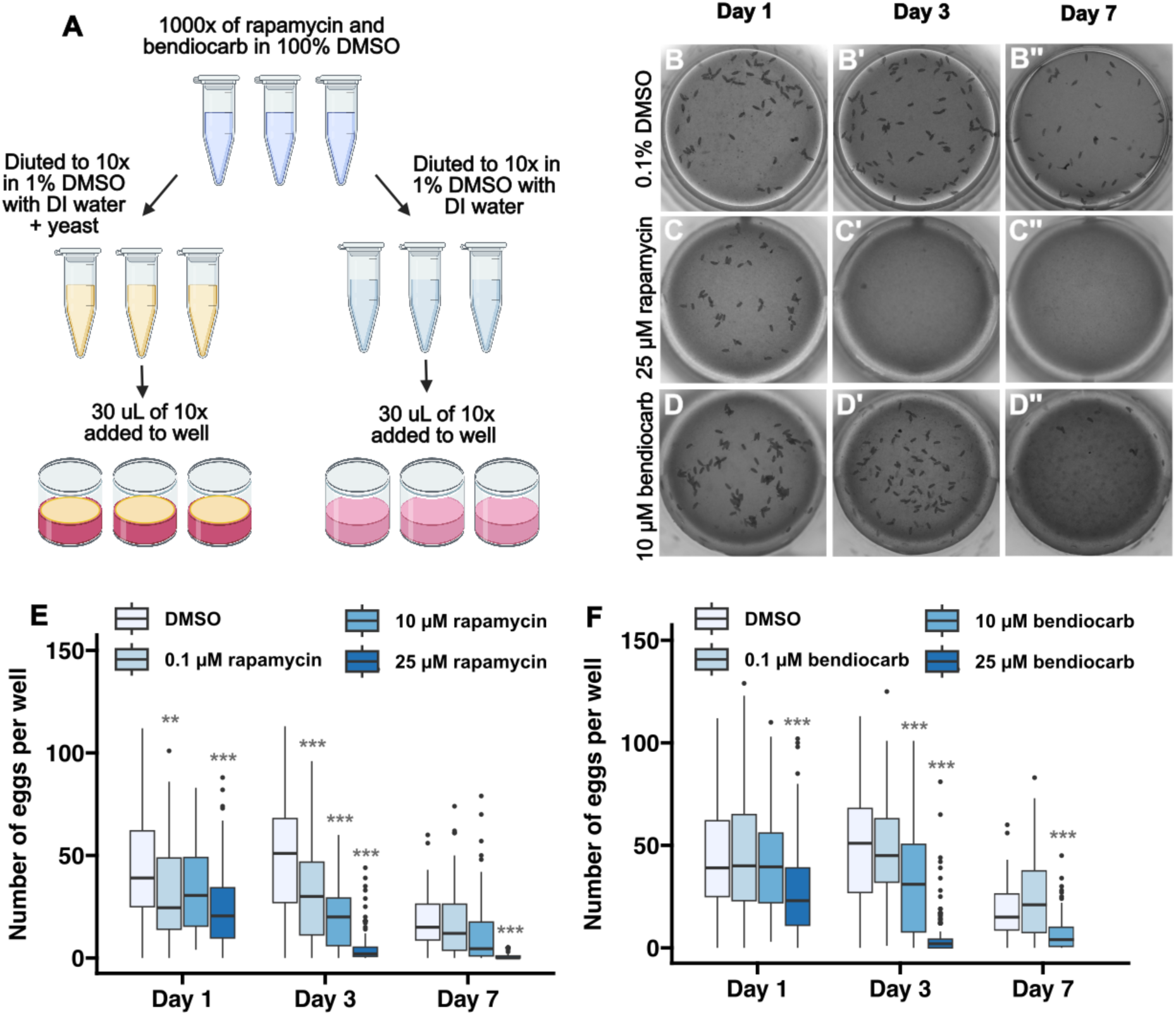
Detection of the impact of dietary exposure to rapamycin and bendiocarb. (A) Diagram showing work flow for fecundity assays in which chemicals are added to the diet. Chemicals are dissolved in 100% DMSO to a 1000x concentration. Then, they are diluted to 10x concentration in 1% DMSO with either water or water plus yeast. Finally, 30 µl of the 10x solution is pipetted onto each well, which contains 300 µl of media, producing a final concentration of 1x chemical in 0.1% DMSO. Using this protocol, flies were exposed to 0.1% DMSO or indicated concentrations of rapamycin or bendiocarb in 0.1% DMSO for 7 days. Egg counts were quantified on days 1, 3, and 7. (B-D) Images of wells at 1, 3 or 7 days after exposure to 0.1% DMSO (B), 25 µM rapamycin (C), or 10 µM bendiocarb (D). Graphs showing the number of eggs laid in the indicated conditions over the 7-day time course. Flies did not survive for 7 days on 25 µM bendiocarb. Asterisks indicate a statistically significant difference compared to the DMSO condition on the same day ** p < 0.01, *** p < 0.001 using pairwise T-tests with a Bonferroni multiple hypothesis test correction.

## Discussion

Here, we present a high-throughput pipeline for measuring Drosophila fecundity that links together multiple innovations. First, the strategy we developed for maintaining robust egg laying in conditions that are compatible with the 48-well format and high-quality imaging of the eggs creates a new opportunity for efficiently testing many different dietary conditions or chemical exposures at a time. The 48-well format substantially increases the efficiency of culturing the flies in different conditions and significantly reduces the cost of media and chemicals, since the volume used for each well is much lower than the volumes used in standard culture vials. Next, we coupled this with custom devices that are low-cost and open source, and we engineered the RoboCam. By integrating consumer-grade 3D printers, cameras, imaging chips, and Raspberry Pi computers, the RoboCam allows the acquisition of high-resolution images in a consistent and accurate way. This utilization of readily available consumer products delivers high precision, reproducibility, and efficiency at low cost, epitomizing the convergence of technology with user accessibility. We implemented a GUI to be used with the Robocam, which provides a user-friendly interface for image collection. In addition, because the RoboCam is a modular platform it is possible to change the different parts for desired functions, allowing for selecting the most suitable camera and lens for different applications and optimizing performance across various tasks. It is noteworthy to underline that the 3D printer within the RoboCam platform retains its original printing function, thus maintaining its dual-purpose capability. From the technical point of view, the protocol’s dedicated operation on the Raspberry Pi ensures the absence of unexpected system updates and the need for a network connection, leading to a more stable and secure operation. Designed for extended use, once that the G-code instructions are programmed through the GUI, the RoboCam can run and record data without human oversight; the recorded data are storage into an external hard drive, guaranteeing quick and efficient data transfers due to immediate access. This arrangement also supports the simultaneous and independent operation of multiple RoboCams, so it can be scaled as needed. The main advancement in the field provided by the Robocam is that it yields reproducible, accurate, and consistent images compared to those obtained with manual methods The RoboCam is thus an efficient and flexible instrument for researchers who aim to harness technology for consistent, high-quality imaging over prolonged periods. Lastly, we have developed an image segmentation process that combines a deep-learning model with additional image segmentation tools to accurately quantify the number of eggs in each image.

Oogenesis in Drosophila is highly conserved, making it a useful model for studies of reproductive health [39]. Indeed 69-78% of genes involved in Drosophila female reproduction have vertebrate orthologs [40], and some aspects, such as meiosis, are particularly well-conserved at both the gene and mechanistic level [41]. The development of this high-throughput method to measure Drosophila fecundity introduces a new tool in the toolbox for these studies. This method is well- suited for toxicology screens to identify compounds in the environment that impact fertility [42,43] and thus aligns well with the need for new approach methodologies (NAMs) that are being adopted by regulatory agencies worldwide [44]. Likewise, this method would also be useful for small molecule screens for therapeutics [45,46] and insect population control [47]. In addition, since oogenesis is highly dependent on and coupled with other organ systems in the body, a high-throughput fecundity screen may be useful for identifying small molecules, dietary conditions or other experimental variables such as age that affect other organs. For example, the rate of oogenesis is regulated by signals from the brain and fat body so changes in fecundity could be used as an assay for RNAi or CRISPR screens that disrupts gene expression in these organs. Moreover, a high-throughput assay for fecundity would be an efficient way to characterize large collections of natural variants [48,49]. Thus, we expect that this new pipeline will have a wide range of applications.

However, because ovarian function depends on other aspects of physiology, an experimental condition that impacts fertility may be operating through any one or more of multiple different mechanisms and follow up experiments would most likely be required to determine why the rate of egg laying is affected. An additional limitation, which may be exacerbated by our strategy of maintaining the flies in small wells on firm media, is that there is substantial variability in the number of eggs laid in each well, which reduces the sensitivity of the assay. We were able to partially counterbalance the variability between individual flies by adding two females into each well and the variability between wells by repeating the same condition in multiple wells. With this approach, the impact of the chemicals we tested was easily detectable, and accepting these limitations in exchange for the ability to screen fecundity in a low-cost, high-throughput manner will likely be a worthwhile tradeoff in many cases. Moreover, our analysis of the downsampled data (**Figure S2**) suggests that similar conclusions can be reached using many fewer wells per condition than we used here. This may be advantageous in situations, such as a large scale screen, where increased in efficiency at the expense of some loss of accuracy is acceptable. Taken together, we expect that this new pipeline will have a wide range of applications in Drosophila research.

## Acknowledgements

We are grateful for the Bloomington *Drosophila* Stock Center for providing the stocks used in this study and for the well-curated and very helpful resource, FlyBase. Diagrams of the workflows show in in Figures 1, 2, and 6 were created with BioRender.com. This work was supported by the following grants from the National Institute of General Medical Sciences GM136348 T.G.N. and GM136348-03S2 to A.G and T.G.N, and T34-GM145400 to S.G. Support for this research was also provided by a CALEPA contract 22-E0032, core center grant P30-ES030284 from the National Institute of Environmental Health Sciences, National Institutes of Health and the Center of Cellular Construction grant STC 1548297 from the National Science Foundation. In addition, this material is based upon work supported by the National Science Foundation under Grant No. DBI-1548297.

## Author contributions

Conceptualization: R.M.E., T.Z., J.C.F., and T.G.N.; Data Curation: A.G. and T.G.N.; Formal Analysis: A.G., A.O., and T.G.N.; Funding Acquisition: R.M.E., S.C., J.C.F., and T.G.N.; Investigation: A.G., S.G., and J.B.D.; Methodology: A.G., A.O., R.M.E., T.Z, J.C.F., and T.G.N.; Project Administration: A.G., R.M.E., S.C., J.C.F., and T.G.N.; Resources: R.M.E., T.Z., J.C.F., and T.G.N.; Software: S.G., A.O., J.B.D., T.Z., and T.G.N.; Supervision: R.M.E., T.Z., S.C., J.C.F., and T.G.N.; Validation: A.G., S.G., A.O., and J.B.D.; Visualization: A.G., S.G., R.M.E., T.Z., S.C., J.C.F., and T.G.N.; Writing – Original Draft Preparation: A.G., A.O., R.M.E, T.Z., S.C., J.C.F. and T.G.N.; Writing – Review and Editing: A.G., A.O., R.M.E., T.Z., S.C., and T.G.N.

---

**Figure S1:**
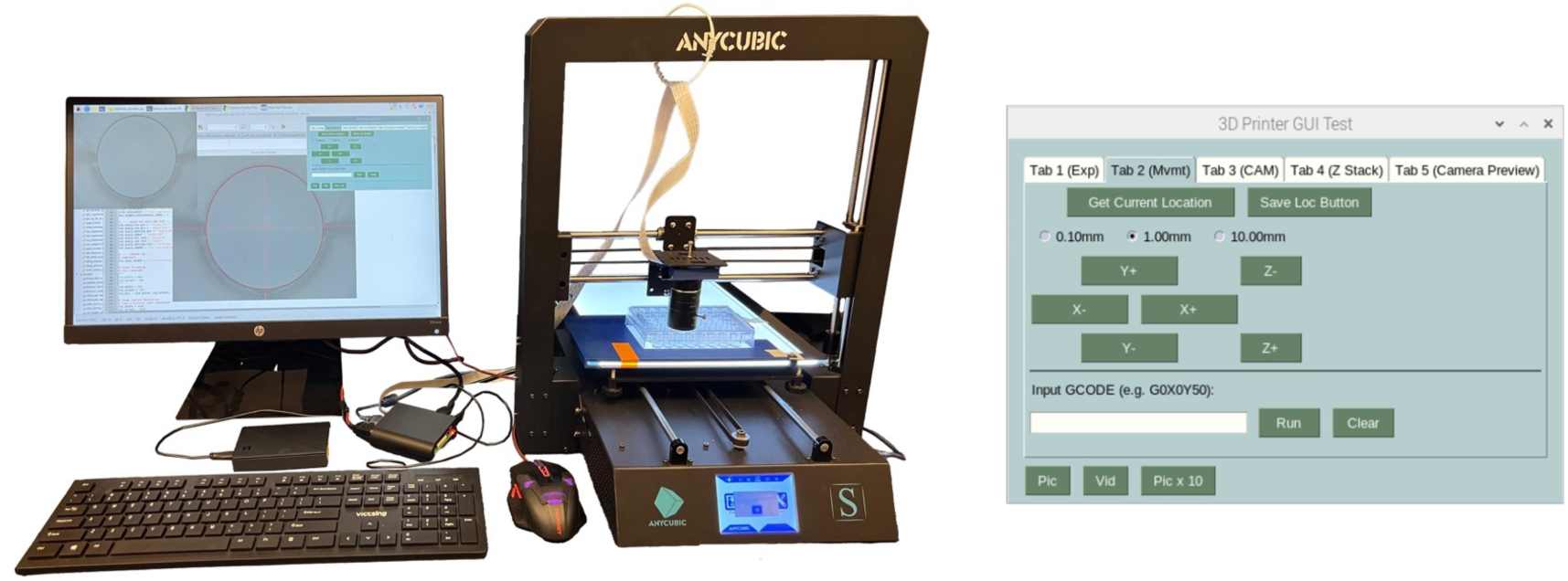
Robocam and graphical user interface. Photograph of the Robocam setup next to a Raspberry Pi, external hard drive, monitor, mouse, and keyboard. Panel on the right shows the graphical user interface of the software that operates the Robocam.

**Figure S2:**
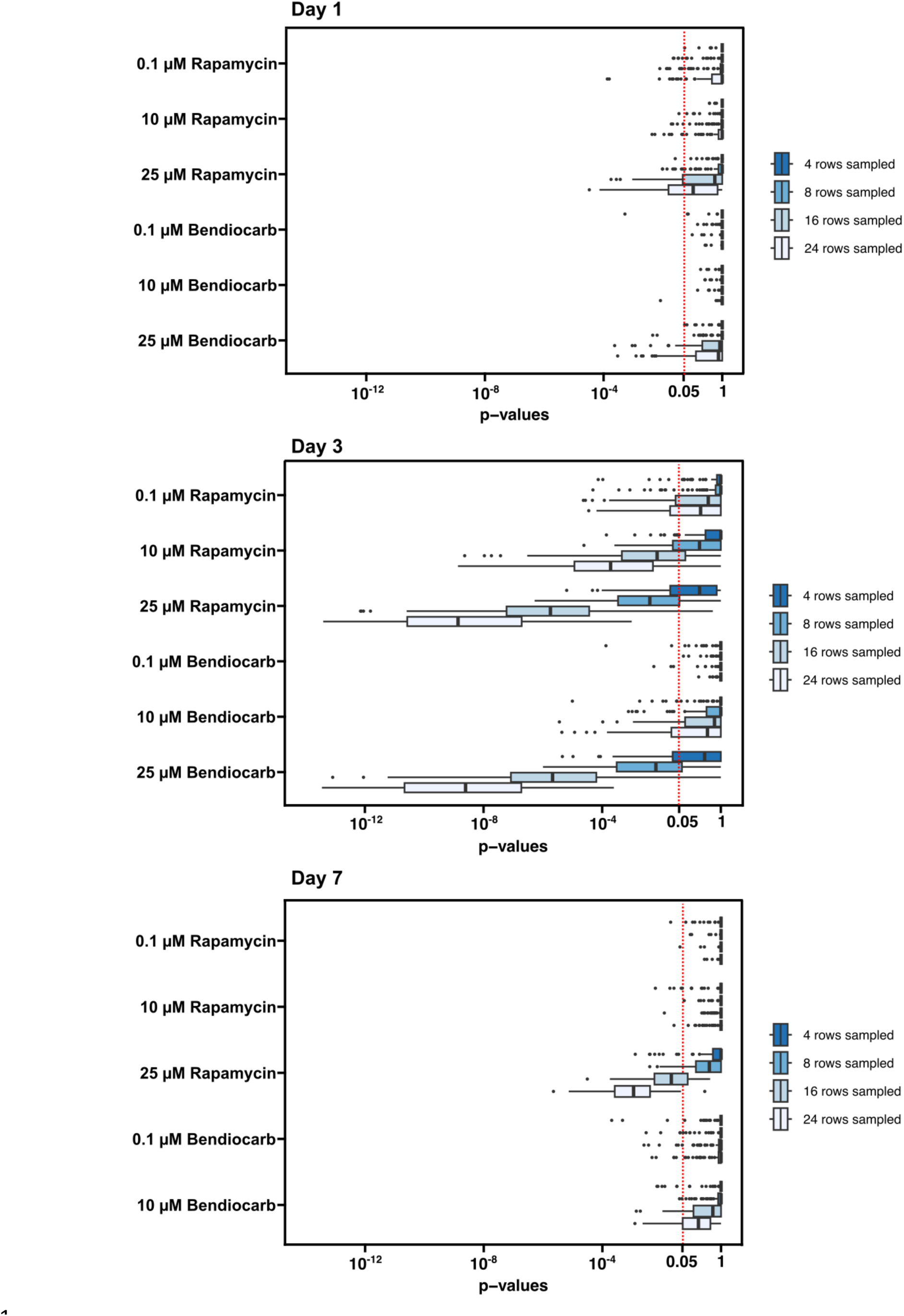
Downsampling of data from dietary exposure to rapamycin and bendiocarb. The p-values from 100 iterations of downsampling the data by randomly selecting 4, 8, 16, or 24 wells from the Day 1, Day 3 and Day 7 datasets are displayed on the graphs. P-values to the left of the red dotted line are below the conventional 0.05 threshold for statistical significance.

